# Developmental emergence of cue-specific fear memory parallels maturation of prefrontal-amygdala circuitry in mice

**DOI:** 10.64898/2026.11.16.767103

**Authors:** Verdion Martina, Zephyr R. Desa, Rodrigo Campos-Cardoso, Terry Lewis, HaoSheng Sun, Kirstie A. Cummings

## Abstract

Fear memories formed in early life are forgotten rapidly, but the circuits that delay stable memory expression are not well defined. The medial prefrontal cortex (mPFC) projects to the basolateral amygdala (BLA), and both regions mature late, yet the behavior, anatomy and physiology of this pathway have rarely been examined together across development. We trained mice in cued auditory fear conditioning at postnatal day (P) 14-15, P20-21 or P59-70 and tested tone-evoked freezing 1, 7 and 14 days later. Early juvenile mice showed no cue-specific freezing at any memory retrieval time points, late juvenile mice showed it at 7 and 14 days but not at 1 day, and adults showed it at all retrieval time points. In separate mice, mPFC axons were low in the BLA at P11 and more abundant at P38 and P90, and light-evoked prelimbic synaptic responses in BLA principal neurons were not detectable at P11 but were robust at P35 and P70. Spontaneous synaptic currents and intrinsic excitability of BLA principal neurons, and excitability of prelimbic principal neurons and interneurons, also changed between P11 and adulthood. These data identify a protracted period of mPFC-BLA circuit maturation that overlaps the emergence of cue-specific fear and define the ages at which candidate mechanisms can be tested.

---

Fear memories formed in infancy are forgotten faster than those formed later in life, a phenomenon known as infantile amnesia that has been observed in every altricial species examined, beginning with rats (Campbell and Campbell 1962; Madsen and Kim 2016; Alberini and Travaglia 2017). In mice, memory for contextual and auditory fear persists for a month only when training occurs at about postnatal day (P) 25 or later; younger mice acquire fear but do not retain it (Akers et al. 2012; Samifanni et al. 2021). The circuits that explain this late onset are not settled. Hippocampal immaturity and high neurogenesis contribute to forgetting of contextual memories (Akers et al. 2014; Travaglia et al. 2016), but they cannot readily explain why hippocampus-independent memories, including fear to a discrete auditory cue, are also forgotten rapidly early in life (Madsen and Kim 2016). Remote memory depends on the medial prefrontal cortex (mPFC) (Frankland and Bontempi 2005), and the prolonged maturation of mPFC circuitry is a leading candidate for why memory persistence emerges late (Klune et al. 2021). A recent preprint links the offset of infantile amnesia for contextual fear to activity-dependent myelination of the prelimbic cortex (Golbabaei et al. 2026), underscoring that prefrontal maturation may be rate-limiting for remote memory.

Within this framework, connectivity between the mPFC and the basolateral amygdala (BLA) is a natural substrate for long-term fear memory. In mice, optogenetic tracing shows that mPFC axons reach other targets, such as the thalamus and striatum, by P10 but are essentially absent from the BLA until P15, when they ramify densely and begin to drive synaptic responses. These synapses then strengthen through P30, when feedforward inhibition transiently peaks (Arruda-Carvalho et al. 2017). This agrees with early tracer work in rats placing the arrival of mPFC axons in the BLA between the second and third postnatal weeks (Bouwmeester et al. 2002a), and mPFC inputs to the basal amygdala are later pruned in late adolescence (Cressman et al. 2010). Anatomical and behavioral studies have largely developed in parallel and the two readouts are rarely compared within one study design. Functional measurements of the circuit, including the intrinsic and synaptic properties of the neurons involved, are also rarely integrated with these readouts.

Here we bridge analyses of behavior, connectivity and function across development. We trained mice at three developmental stages (P14-15, P20-21 and P59-70; early juvenile, late juvenile and adult in the scheme of Madsen and Kim (Madsen and Kim 2016)) in a cued auditory fear conditioning paradigm and tested cue-evoked freezing 1, 7 and 14 days later. In parallel, in separate mice, we traced mPFC output to the BLA with an anterograde virus at P11 (late infancy), P38 (adolescence) and P90 (adulthood), and used whole-cell recordings to measure synaptic input from the prelimbic cortex onto BLA principal neurons and the intrinsic excitability of BLA and prelimbic neurons across development. We find that cue-specific fear emerges in stages, being absent in early juvenile mice, begins to emerge in late juvenile mice and strengthens into adulthood. We also find that mPFC axons and functional prelimbic input to the BLA are low or undetectable at P11 and present by P35-38. Together these data define a window of prefrontal-amygdala circuit maturation that overlaps the emergence of cue-specific fear, and they set the stage for testing the molecular programs that build this circuit.

## RESULTS

### Cue-specific auditory fear emerges across development

To determine whether the ability to express auditory fear memory differs across development, we used cued auditory fear conditioning (Fig. 1A). Mice were trained at P14-15 (early juvenile), P20-21 (late juvenile) or P59-70 (adult) (n = 14, 16, 16; both sexes) with six tone-shock pairings. Cue-evoked freezing was then assessed in a novel context 1 d (24 hours) after training (recent memory) and 7 and 14 d after training (remote memory).

**Figure 1.**
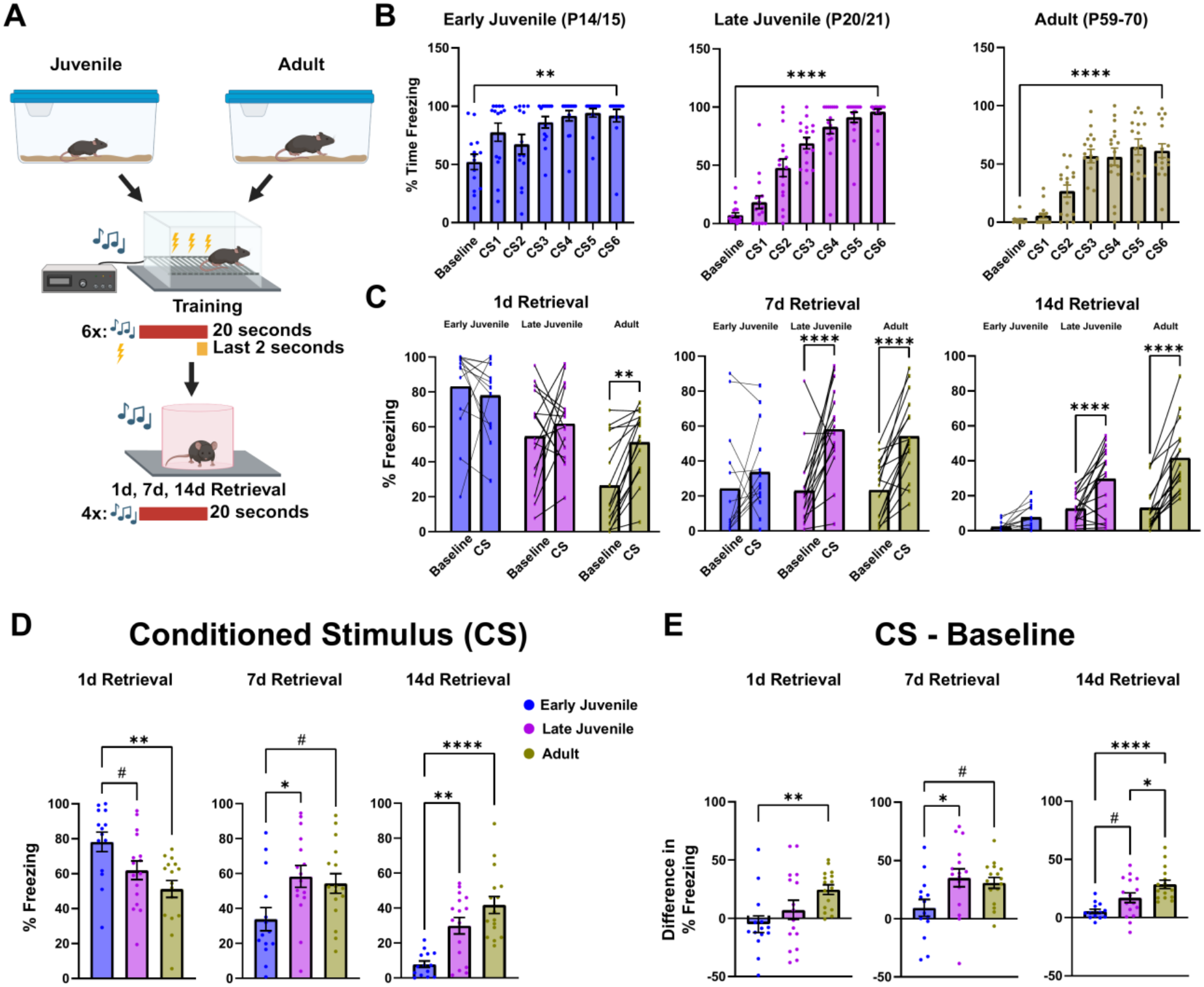
Cue-specific auditory fear emerges across development. **(A)** Cued auditory fear conditioning. Mice were trained at P14-15 (early juvenile), P20-21 (late juvenile) or P59-70 (adult) with six pairings of a tone (CS; 2 kHz, 80 dB, 20 s) that co-terminated with a footshock (US; 0.6 mA, 2 s). Memory was tested in a novel context with four tones 1, 7 and 14 d later. **(B)** Percent time freezing during the pre-tone baseline and each CS presentation (CS1-CS6) during training, analyzed within each age group by one-way repeated-measures ANOVA (Geisser-Greenhouse correction) with Dunnett’s multiple comparisons test against baseline. Early juvenile (n = 14): F(3.659, 47.56) = 6.901, P = 0.0003. Baseline vs. CS1, P = 0.0475; CS2, P = 0.4651; CS3, P = 0.0184; CS4, P = 0.0027; CS5, P < 0.0001; CS6, P = 0.0032. Late juvenile (n = 16): F(3.521, 52.81) = 55.93, P < 0.0001. CS1, P = 0.2149; CS2, P = 0.0002; CS3-CS6, P < 0.0001. Adult (n = 16): F(3.231, 48.46) = 29.44, P < 0.0001. CS1, P = 0.1719; CS2, P = 0.0008; CS3-CS6, P < 0.0001. Asterisks indicate baseline vs. CS6. **(C)** Percent time freezing during the pre-tone baseline and the mean of the four CS presentations at each memory retrieval time point; lines connect individual mice. Each retrieval time point was analyzed by two-way repeated-measures ANOVA (stimulus, matched; developmental stage), followed by Šídák’s multiple comparisons test of baseline vs. CS within each stage (n = 14 early juvenile, 16 late juvenile, 16 adult). 1 d: Stage × stimulus interaction, F(2,43) = 4.872, P = 0.0124. Stimulus, F(1,43) = 5.297, P = 0.0263. Stage, F(2,43) = 18.04, P < 0.0001. Baseline vs. CS: early juvenile, P = 0.8565; late juvenile, P = 0.6272; adult, P = 0.0015. 7 d: Interaction, F(2,43) = 4.194, P = 0.0217. Stimulus, F(1,43) = 43.57, P < 0.0001. Stage, F(2,43) = 1.403, P = 0.2568. Baseline vs. CS: early juvenile, P = 0.4467; late juvenile, P < 0.0001; adult, P < 0.0001. 14 d: Interaction, F(2,43) = 11.34, P = 0.0001. Stimulus, F(1,43) = 75.45, P < 0.0001. Stage, F(2,43) = 14.58, P < 0.0001. Baseline vs. CS: early juvenile, P = 0.3423; late juvenile, P < 0.0001; adult, P < 0.0001. **(D)** Percent time freezing during the CS at each retrieval time point, compared across developmental stages by one-way ANOVA with Tukey’s multiple comparisons test, run separately for each retrieval time point. 1 d: F(2,43) = 6.577, P = 0.0032. Early juvenile vs. late juvenile, P = 0.0856; early juvenile vs. adult, P = 0.0022; late juvenile vs. adult, P = 0.3098. 7 d: F(2,43) = 4.338, P = 0.0192. Early vs. late juvenile, P = 0.0219; early juvenile vs. adult, P = 0.0635; late juvenile vs. adult, P = 0.8873. 14 d: F(2,43) = 16.08, P < 0.0001. Early vs. late juvenile, P = 0.0021; early juvenile vs. adult, P < 0.0001; late juvenile vs. adult, P = 0.1129. **(E)** Difference in freezing between CS and baseline (CS − baseline) at each retrieval time point, analyzed as in D. 1 d: F(2,43) = 4.872, P = 0.0124; early vs. late juvenile, P = 0.4187; early juvenile vs. adult, P = 0.0096; late juvenile vs. adult, P = 0.1562. 7 d: F(2,43) = 4.194, P = 0.0217; early vs. late juvenile, P = 0.0241; early juvenile vs. adult, P = 0.0716; late juvenile vs. adult, P = 0.8791. 14 d: F(2,43) = 11.34, P = 0.0001; early vs. late juvenile, P = 0.0554; early juvenile vs. adult, P < 0.0001; late juvenile vs. adult, P = 0.0465. Bars show mean ± SEM; dots show individual mice. #0.05<P<0.1; *P < 0.05, **P < 0.01, ***P < 0.001, ****P < 0.0001.

During training, freezing increased across pairings in all groups (one-way repeated-measures ANOVA: early juvenile, F(3.659, 47.56) = 6.901, P = 0.0003; late juvenile, F(3.521, 52.81) = 55.93, P < 0.0001; adult, F(3.231, 48.46) = 29.44, P < 0.0001; Fig. 1B). Early juvenile mice, however, were immobile for roughly half of the pre-tone baseline, before any shock (~52%), whereas late juvenile and adult mice rarely were (~7% and ~2%). Because this immobility cannot be attributed to learned fear, we assessed memory at retrieval by comparing freezing during the tone (CS) with the preceding baseline in each mouse (Fig. 1C), by freezing at CS alone (Fig. 1D), and by the CS − baseline difference (Fig. 1E).

At 1 d after training, there was a significant age × stimulus interaction (two-way repeated-measures ANOVA, F(2,43) = 4.872, P = 0.0124). Adults froze significantly more during the tone than at baseline (Šídák’s test, P = 0.0015), whereas neither early (P = 0.8565) nor late juveniles did (P = 0.6272; Fig. 1C), and the CS − baseline difference was smaller in early juveniles than in adults (Tukey’s test, P = 0.0096; Fig. 1E). Pre-tone freezing in the novel context was high in juveniles (~83% and ~54%) relative to adults (~26%). In late juveniles, this was well above their pre-shock baseline (~7%), consistent with generalization of conditioned fear to the novel context. In early juveniles, pre-shock immobility (~50%) confounds this interpretation.

At 7 d after training, the interaction remained significant (F(2,43) = 4.194, P = 0.0217): late juvenile and adult mice showed cue-specific freezing (both P < 0.0001), whereas early juveniles did not (P = 0.4467; Fig. 1C), and the CS − baseline difference was smaller in early than in late juveniles (P = 0.0241; Fig. 1E) and adults (P= 0.0716). At 14 d, the interaction was strongest (F(2,43) = 11.34, P = 0.0001): late juveniles and adults still froze more during the tone (both P < 0.0001), with a larger increase in adults than in early (P < 0.0001) or late juveniles (P = 0.0465; Fig. 1C,E), and early juveniles froze very little in either baseline or CS conditions (~2% and ~8%) and showed no significant increase (P = 0.3423). Thus, early juvenile mice showed no cue-specific freezing at any retrieval time point, late juvenile mice showed cue-specific freezing at 7 and 14 d but not 1 d, and adults showed it at every retrieval time point.

### mPFC projections to the BLA are low at P11 and present by P38

Because fear learning and expression depends on prefrontal-amygdala circuitry (Corcoran and Quirk 2007; Arruda-Carvalho and Clem 2014; Tovote et al. 2015), we asked whether mPFC projections to the BLA differ across the ages at which cue-specific fear differs. We bilaterally injected an AAV1 encoding channelrhodopsin-2 fused to eYFP under the CaMKIIα promoter (AAV1-CaMKII-ChR2-eYFP) into the mPFC at P1, P28 or P80 and examined the brains 10 d later, at P11, P38 and P90 (Fig. 2A; n = 4, 4 and 5 mice, respectively; both sexes). Injection volume and coordinates were adjusted for brain size at each age. The percentage of mPFC area expressing eYFP did not differ significantly across ages (one-way ANOVA, F(2,10) = 3.078, P = 0.0909; Tukey’s test: P11 vs. P38, P = 0.3292; P11 vs. P90, P = 0.6630; P38 vs. P90, P = 0.0778; Fig. 2B), and expression was centered on the prelimbic cortex at every age. eYFP-labeled fibers were also present in the mediodorsal and ventromedial thalamus, which receive mPFC projections, at all ages, indicating successful expression and axonal transport at P11 (Fig. 2C).

**Figure 2.**
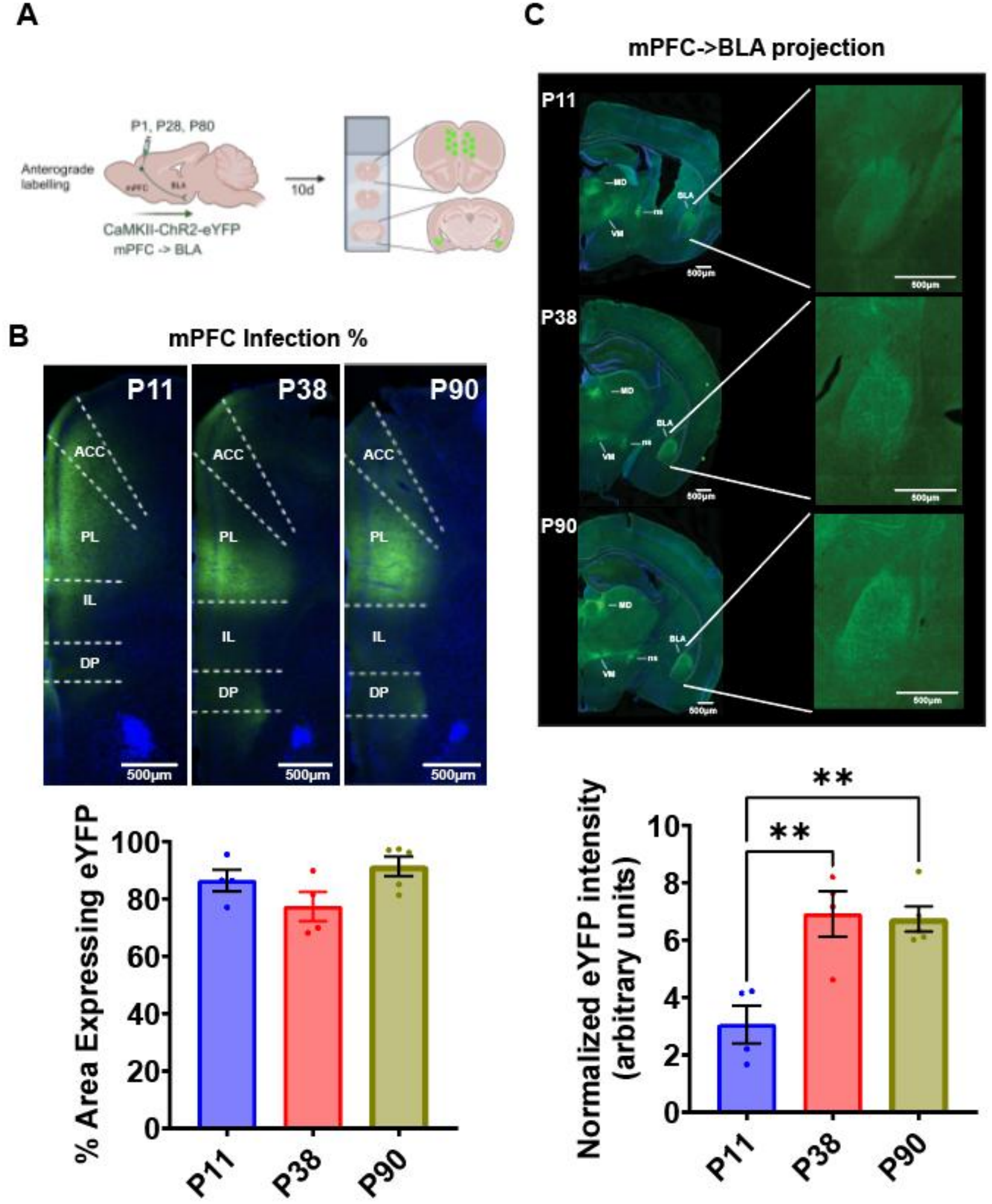
mPFC projections to the BLA are low at P11 and present by P38. **(A)** Experimental design. AAV1-CaMKIIα-hChR2(H134R)-eYFP was injected bilaterally into the mPFC at P1, P28 or P80, and brains were collected 10 d later (P11, P38, P90), sectioned and imaged. **(B)** Top, representative sections showing eYFP (green) and DAPI (blue) in the mPFC at each age (ACC, anterior cingulate; PL, prelimbic; IL, infralimbic; DP, dorsal peduncular). Scale bars, 500 μm. Bottom, percentage of mPFC area expressing eYFP. Infection did not differ significantly across ages (one-way ANOVA, F(2,10) = 3.078, P = 0.0909; Tukey’s multiple comparisons test: P11 vs. P38, P = 0.3292; P11 vs. P90, P = 0.6630; P38 vs. P90, P = 0.0778). **(C)** Top, representative sections showing eYFP in the mediodorsal (MD), ventromedial (VM) thalamus, * (mPFC fiber tracks) and the BLA at each age, with higher-magnification insets of the BLA; imaging and display settings were identical across ages. Scale bars, 500 μm. Bottom, BLA eYFP intensity normalized to the percentage of mPFC area expressing eYFP in the same animal (B). Normalized signal differed across ages (one-way ANOVA, F(2,10) = 11.82, P = 0.0023; Tukey’s test: P11 vs. P38, P = 0.0045; P11 vs. P90, P = 0.0043; P38 vs. P90, P = 0.9785). Bars show mean ± SEM; dots show individual mice (n = 4 P11, 4 P38, 5 P90; both sexes). **P < 0.01.

To account for differences in viral transduction between animals, we normalized BLA eYFP intensity to the percentage of mPFC area expressing eYFP in the same animal. Normalized BLA signal, measured under identical acquisition settings, differed across ages (one-way ANOVA, F(2,10) = 11.82, P = 0.0023; Fig. 2C). It was lower at P11 than at P38 (Tukey’s test, P = 0.0045) or P90 (P = 0.0043) and did not differ between P38 and P90 (P = 0.9785). These data indicate that mPFC axons innervate the BLA substantially only after P11, and that this innervation is present by P38 and maintained into adulthood. Late juvenile mice (P20-21), the youngest group that expressed cue-specific fear, fall between our P11 and P38 tracing ages.

### Prelimbic input onto BLA principal neurons is not detectable at P11 and increases across development

Our circuit tracing (Fig. 2) revealed developmentally regulated changes in the projection from the mPFC to the BLA. We therefore asked how synaptic input onto BLA principal neurons (PNs) changes across development. We recorded miniature excitatory (mEPSC) and inhibitory (mIPSC) postsynaptic currents in the presence of 1 µM tetrodotoxin from BLA PNs, identified by morphology, in P11, P32-36 (hereafter P35) and P70 mice (Fig. 3A-D). The mEPSC interevent interval was longer at P11 than at P70 (one-way ANOVA, F(2,31) = 4.39, P = 0.021; Tukey’s test, P < 0.05), indicating a lower event frequency early in development, with no group differences in amplitude (F(2,31) = 2.06, P = 0.15; Fig. 3B). The mIPSC interevent interval was also longer at P11 than at P70 (Kruskal-Wallis, χ^2^(2) = 12.6, P = 0.002; Dunn’s test, P < 0.01), with a trend between P11 and P35 (P = 0.08), and mIPSC amplitude was larger at P35 than at P11 (F(2,30) = 3.70, P = 0.03; Tukey’s test, P < 0.05; Fig. 3D). These results are consistent with both long-range projections and local microcircuitry undergoing significant change across these ages.

**Figure 3.**
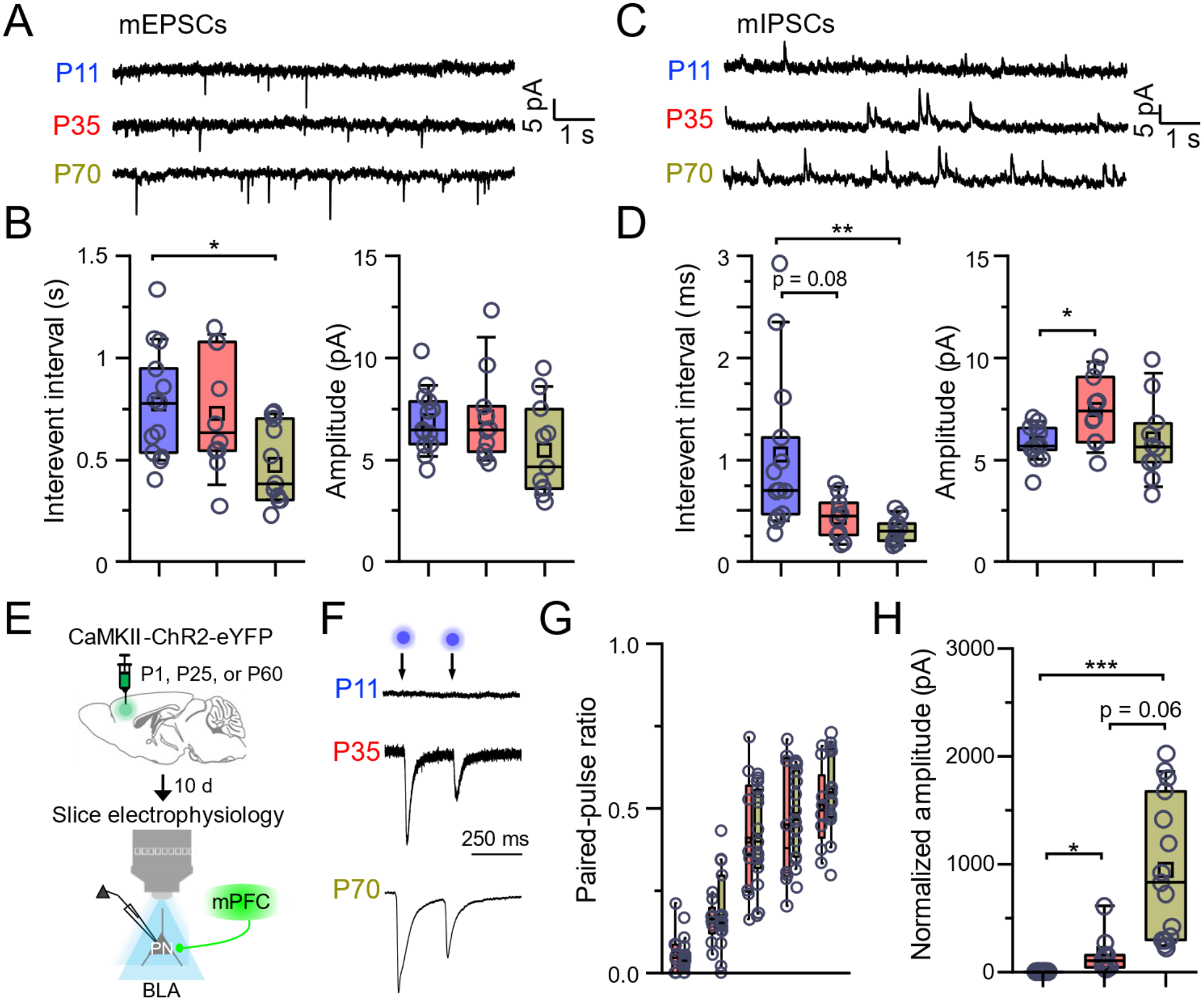
Synaptic measurements from basolateral amygdala principal neurons in P11, P32-36 and P70 mice. **(A)** Example miniature excitatory postsynaptic current (mEPSC) recordings from P11 (n = 13 cells from 2 mice, 1 female and 1 male), P32-36 (n = 10 cells from 3 mice, 1 male and 2 females) and P70 (n = 10-11 cells from 3 mice, 1 female and 2 males) mice. **(B)** mEPSC interevent interval (one-way ANOVA, F(2,31) = 4.39, P = 0.021; * P < 0.05, Tukey’s test) and amplitude (F(2,31) = 2.06, P = 0.15). **(C)** Example miniature inhibitory postsynaptic current (mIPSC) recordings from the same cells. **(D)** mIPSC interevent interval (Kruskal-Wallis, χ^2^(2) = 12.6, P = 0.002; ** P < 0.01, Dunn’s test; P11 vs. P32-36, P = 0.08) and amplitude (one-way ANOVA, F(2,30) = 3.70, P = 0.03; * P < 0.05, Tukey’s test). **(E)** Strategy for light-evoked currents: AAV1-CaMKII-ChR2-eYFP was injected into the PL at P1, P25 or P60 and slices were prepared 10 d later. Recordings were made at P11 (n = 14 cells from 4 mice, 2 females and 2 males), P35 (n = 8-9 cells from 3 mice, 2 males and 1 female) and P70 (n = 15 cells from 4 mice, 3 males and 1 female). **(F)** Representative monosynaptic light-evoked paired-pulse responses from PL axons onto BLA PNs at a 250 ms delay interval. Response amplitudes are normalized to amplitude of first response for P35 and P70. **(G)** Paired-pulse ratios for P35 and P70 mice across interstimulus intervals (interaction between age and ratio: two-way repeated-measures ANOVA, F(4,28) = 0.51, P = 0.73). **(H)** BLA PN current amplitudes normalized to the area of ChR2-transduced PL (Kruskal-Wallis, χ^2^(2) = 35.6, P = 0.0001; * P < 0.05, *** P < 0.001, Dunn’s test; P35 vs. P70, P = 0.06). Boxplots show the median (line), mean (square), quartiles and 10-90% range (whiskers); open circles show individual cells.

Given the importance of the prelimbic cortex (PL), the BLA and their connectivity for fear memory (Corcoran and Quirk 2007), we next used optogenetics-assisted electrophysiology to isolate input from PL axon terminals onto BLA PNs. Mice received AAV1-CaMKII-ChR2-eYFP in the PL at P1, P25 or P60, and acute BLA slices were prepared 10 d later, at P11, P35 and P70 (Fig. 3E). Light-evoked monosynaptic EPSCs were recorded in the presence of 1 µM tetrodotoxin and 100 µM 4-aminopyridine. Consistent with our tracing results (Fig. 2C), we could not detect light-evoked responses from PL axons onto BLA PNs at P11 (n = 14 cells from 4 mice; Fig. 3F,H), in line with the lower mEPSC frequency at that age (Fig. 3B). At P35 and P70, we recorded robust light-evoked currents in all BLA PNs patched (n = 8 cells from 3 mice and n = 15 cells from 4 mice, respectively). Paired-pulse ratios did not differ between P35 and P70 (two-way repeated-measures ANOVA, F(4,28) = 0.51, P = 0.73; Fig. 3G), consistent with unchanged glutamate release probability across this period. Current amplitudes normalized to the area of ChR2-transduced PL, however, differed across ages (Kruskal-Wallis, χ^2^(2) = 35.6, P = 0.0001; Fig. 3H): amplitudes were larger at P35 than at P11 (Dunn’s test, P < 0.05) and larger at P70 than at P11 (P < 0.001), with a trend toward larger amplitudes at P70 than at P35 (P = 0.06). These results may reflect developmental changes in the number of functional PL inputs rather than in release probability.

### Intrinsic excitability of BLA principal neurons decreases across development

As synaptic connections reach their targets and mature, alterations in intrinsic excitability are also prominent (Ehrlich and Lauri 2026). Because excitability is a critical determinant of memory encoding (Yiu et al. 2014; Abatis et al. 2024), we asked how the intrinsic properties of BLA PNs change across development. We performed whole-cell recordings from BLA PNs in acute slices from P11, P32-36 and P70 mice, identified by morphology and confirmed by membrane properties and spike waveform, in the presence of blockers of synaptic transmission (10 µM CNQX, 75 µM AP5 and 100 µM picrotoxin). BLA PNs from P11 mice fired more action potentials at the same current injections than PNs from P32-36 and P70 mice (two-way repeated-measures ANOVA, age × current interaction, F(38,342) = 1.99, P = 7.7 × 10−^4^; Fig. 4A,B). Rheobase was lower (one-way ANOVA, F(2,42) = 4.66, P = 0.015; Tukey’s test, P11 vs. P32-36 and P11 vs. P70, both P < 0.05; Fig. 4C) and membrane resistance was higher (Kruskal-Wallis, χ^2^(2) = 19.97, P = 0.0001; Dunn’s test, both P < 0.001; Fig. 4D) in P11 neurons. Resting membrane potential (F(2,42) = 0.8, P = 0.46) and firing threshold (F(2,42) = 0.69, P = 0.51) did not differ across ages (Fig. 4E,F). Capacitance increased from P11 to P32-36 and P70 (Kruskal-Wallis, χ^2^(2) = 14.78, P = 6.15 × 10−^4^; Dunn’s test, P11 vs. P32-36, P < 0.05; P11 vs. P70, P < 0.001; Fig. 4G), likely reflecting the growth of dendritic arbors and soma size across these ages (Ryan et al. 2016). Overall, the age-dependent changes in firing frequency, rheobase and membrane resistance reflect a decrease in BLA PN intrinsic excitability between early and late developmental periods.

**Figure 4.**
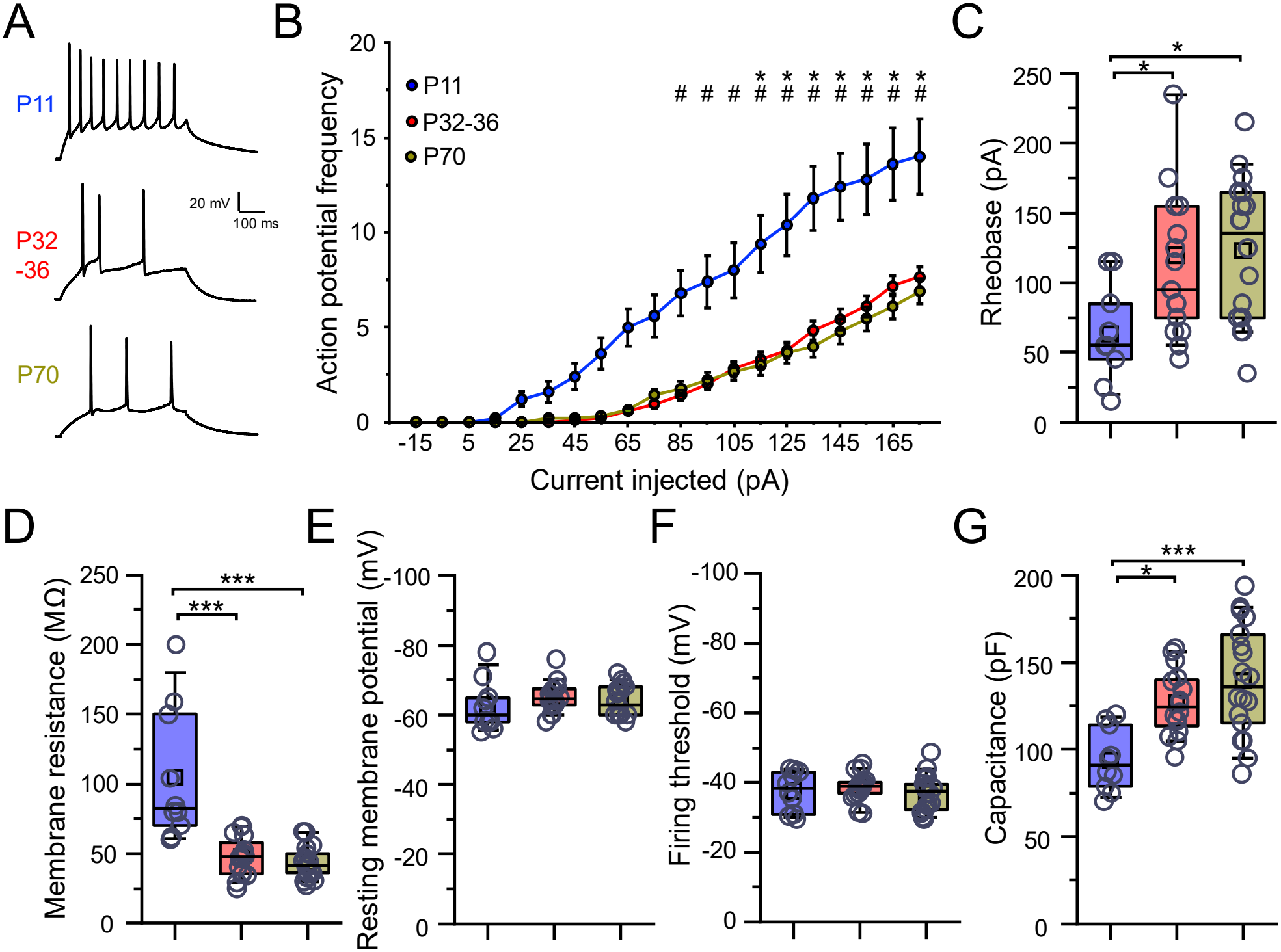
Intrinsic excitability of basolateral amygdala principal neurons in P11, P32-36 and P70 mice. **(A)** Example action potentials elicited at 125 pA current injection in P11 (n = 10 cells from 2 mice, both female), P32-36 (n = 17 cells from 4 mice, 3 females and 1 male) and P70 (n = 18 cells from 3 mice, 1 female and 2 males) mice. **(B)** Action potential input-output. Significant age × current interaction: F(38,342) = 1.99, P = 7.7 × 10−^4^, two-way repeated-measures ANOVA. * P < 0.05 for P70 vs. P11; # P < 0.05 for P32-36 vs. P11, Tukey’s test. Quantification across ages for **(C)** rheobase (F(2,42) = 4.66, P = 0.015, one-way ANOVA; * P < 0.05, Tukey’s test), **(D)** membrane resistance (χ^2^(2) = 19.97, P = 0.0001, Kruskal-Wallis; *** P < 0.001, Dunn’s test), **(E)** resting membrane potential (F(2,42) = 0.8, P = 0.46, one-way ANOVA), **(F)** firing threshold (F(2,42) = 0.69, P = 0.51, one-way ANOVA) and **(G)** capacitance (χ^2^(2) = 14.78, P = 6.15 × 10−^4^, Kruskal-Wallis; * P < 0.05, *** P < 0.001, Dunn’s test). Boxplots show the median (line), mean (square), quartiles and 10-90% range (whiskers); open circles show individual cells.

### Prelimbic principal neurons and interneurons show opposite patterns of intrinsic excitability across development

We next made the same measurements in the PL at P10-11 and P70. In addition to PNs, we recorded from somatostatin (SST-INs) and parvalbumin (PV-INs) interneurons, given their involvement in fear memory (Courtin et al. 2014; Cummings and Clem 2020; Cummings et al. 2022). SST-Flp/Ai65F or PV-Cre/Ai9 transgenic mice were used to visualize interneurons via tdTomato fluorescence. As in the BLA, PL PNs from P70 mice fired fewer action potentials in response to increasing current than PNs from P10-11 mice (age × current interaction, F(10,160) = 24.5, P = 0.0001; Fig. 5A,B). They also had a higher rheobase (Mann-Whitney U = 22.5, P = 1.11 × 10−^4^; Fig. 5C) and a more negative resting membrane potential (U = 267, P = 0.0001; Fig. 5D). Interneurons showed the opposite pattern for firing: SST-INs and PV-INs from P70 mice fired more at higher current injections than those from P10-11 mice (age × current interaction: SST-INs, F(10,120) = 11.4, P = 0.0001; PV-INs, F(10,40) = 11.1, P = 0.0001; Fig. 5E,F,I,J). Rheobase did not differ in SST-INs (U = 132, P = 0.26) and trended higher in PV-INs at P70 (U = 8, P = 0.058; Fig. 5G,K). Resting membrane potential decreased in SST-INs across development (t(37) = −2.33, P = 0.025) but not in P-INs (U = 14.5, P = 0.31; Fig. 5H,L). Thus, PL PNs decrease in excitability across development, as BLA PNs do, whereas SST-INs and PV-INs exhibit enhancements in excitability, supporting the idea that the mPFC microcircuit changes substantially across these ages.

**Figure 5.**
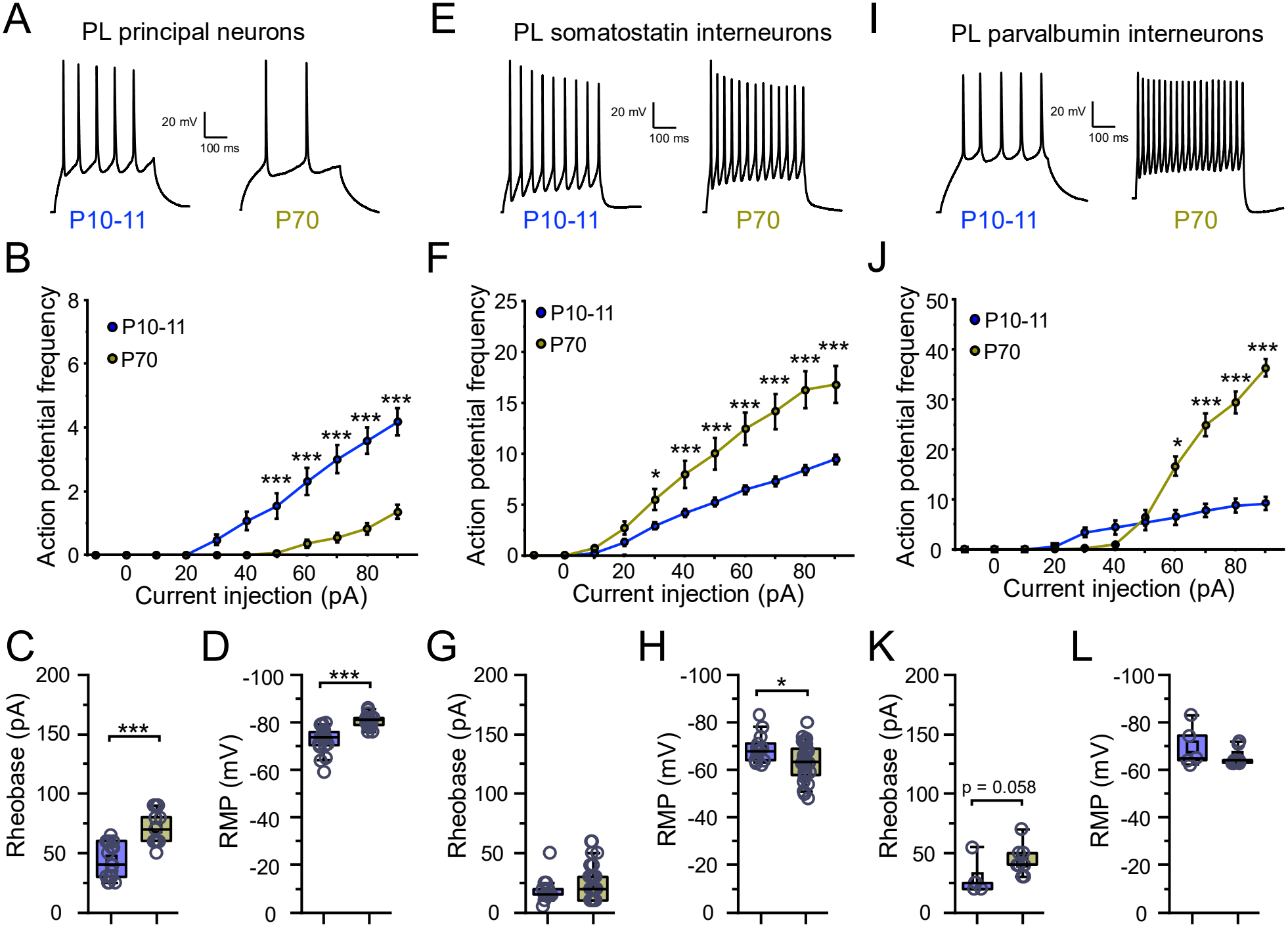
Intrinsic excitability of prelimbic principal neurons, somatostatin interneurons and parvalbumin interneurons in P11 and P70 mice. **(A)** Example action potentials elicited in PNs at 90 pA in P11 (n = 17 cells from 4 mice, 2 males and 2 females) and P70 (n = 17 cells from 3 mice, 2 males and 1 female) mice. **(B)** PN action potential input-output; age × current interaction, F(10,160) = 24.5, P = 0.0001, two-way repeated-measures ANOVA; *** P < 0.001 for P70 vs. P11, Tukey’s test. PN **(C)** rheobase (Mann-Whitney U = 22.5, P = 1.11 × 10−^4^) and **(D)** resting membrane potential (U = 267, P = 0.0001). **(E)** Example action potentials in SST-INs at 90 pA in P11 (n = 13 cells from 2 mice, 1 male and 1 female) and P70 (n = 26 cells from 4 mice, 2 male and 2 female) mice. **(F)** SST-IN input-output; age × current interaction, F(10,120) = 11.4, P = 0.0001; * P < 0.05, *** P < 0.001 for P70 vs. P11, Tukey’s test. SST-IN **(G)** rheobase (U = 132, P = 0.26) and **(H)** resting membrane potential (t(37) = −2.33, P = 0.025, two-tailed unpaired t-test). **(I)** Example action potentials in PV-INs at 90 pA in P11 (n = 5 cells from 2 mice, 1 male and 1 female) and P70 (n = 9 cells from 2 mice, 1 male and 1 female) mice. **(J)** PV-IN input-output; age × current interaction, F(10,40) = 11.1, P = 0.0001; * P < 0.05, *** P < 0.001 for P70 vs. P11, Tukey’s test. PV-IN **(K)** rheobase (U = 8, P = 0.058) and **(L)** resting membrane potential (U = 14.5, P = 0.31). Boxplots show the median (line), mean (square), quartiles and 10-90% range (whiskers); open circles show individual cells.

## DISCUSSION

Early juvenile mice did not express cue-specific fear at any memory retrieval time points, late juvenile mice expressed it from 7 d, and adults at all time points. Over the same developmental period, mPFC axons in the BLA, functional prelimbic input onto BLA principal neurons, and the synaptic and intrinsic properties of BLA and prelimbic neurons changed substantially. Together, these data define a period of prefrontal-amygdala circuit maturation that overlaps the emergence of cue-specific fear.

### Behavior

Our finding that early-trained mice fail to express cue-specific fear agrees with the emergence of persistent auditory and contextual fear memory at around P25 (Samifanni et al. 2021), and with the view that infantile amnesia reflects immature consolidation or retrieval systems rather than a failure to encode. Juvenile rodents learn at rates comparable to older animals (Fig. 1B) but forget faster, and stronger training does not rescue retention (Madsen and Kim 2016; Alberini and Travaglia 2017). Previous studies (Samifanni et al. 2021) also found that remote contextual fear recruits prelimbic c-Fos at P30 but not at P15, tying the onset of memory persistence to prefrontal engagement, and a recent preprint ties the offset of infantile amnesia to prelimbic myelination (Golbabaei et al. 2026). One feature of our data deserves care: early juvenile mice exhibited substantial immobility before the first tone during training, so freezing in this group cannot be read as a pure index of learned fear. At 1 d, early juveniles were immobile at high levels during both baseline and tone with no cue-evoked increase, and at 7 and 14 d they froze little. We therefore describe the early juvenile phenotype as an absence of cue-specific freezing at every retrieval time points, rather than as a selective loss of remote memory. Late juvenile mice showed a different pattern: no cue-specific freezing at 1 d, when pre-tone freezing in the novel context was high, and robust cue-specific freezing at 7 and 14 d as pre-tone freezing declined. Whether high pre-tone freezing at 1 d reflects generalization of conditioned fear that masks cue-specific responding will need no-shock and unpaired controls. Our late juvenile result also differs from that of Samifanni et al. (Samifanni et al. 2021), in whom mice trained at P21 showed minimal freezing 30 d later. The shorter interval in our study (14 d) and differences in tone and shock parameters may all contribute.

Because the same mice were tested at 1, 7 and 14 d, earlier retrieval sessions could influence later ones. Our retrieval sessions were brief (four CS presentations) and are unlikely to constitute extinction training. Repeated retrieval can, however, destabilize and update memories (Misanin et al. 1968; Nader et al. 2000; Eisenberg et al. 2003; Dudai 2006; Alberini 2011), which could alter memory strength. We observed freezing levels similar to those in our previous studies, in which memory strength was examined at these time points in separate cohorts of mice (Cummings and Clem 2020; Cummings et al. 2022; Campos-Cardoso et al. 2024; Campos Cardoso et al. 2026).

### Anatomy

The low mPFC signal in the BLA at P11 and its presence at P38 and P90 reproduces the late arrival of this projection seen with the same viral strategy, in which mPFC axons reached other targets by P10 but entered the BLA between P10 and P15 and no light-evoked synaptic responses were recorded at P10 (Arruda-Carvalho et al. 2017), and with classical tracing in rats (Bouwmeester et al. 2002a). Labeling in thalamic and other control regions argues against a general expression or transport deficit at P11. The same earlier work found dense mPFC fibers in the BLA by P15 and synaptic strengthening between P15 and P30 (Arruda-Carvalho et al. 2017). If so, mPFC axons are likely present in the BLA of early juvenile mice trained at P14-15 and tested through about P29, when they still lack cue-specific fear, so axonal arrival alone cannot explain our behavioral result. Maturation of synaptic strength or inhibition, or of other circuits, may. We did not examine the BLA-to-mPFC projection, which arrives earlier than the mPFC output in rats (Bouwmeester et al. 2002a; Bouwmeester et al. 2002b).

### Electrophysiology

In line with our circuit tracing, we were unable to detect functional PL axon terminals in the BLA of P11 mice, consistent with the idea that these projections have not yet reached the BLA at this age. This agrees with previous work (Arruda-Carvalho et al. 2017) and correlates with increases in glutamatergic transmission onto BLA PNs from P11 to P70. We also observed increases in GABAergic currents onto BLA PNs across development. In rats, parvalbumin-expressing interneurons (PV-INs) begin integrating into the BLA circuit around P10 and complete this process around P30 (Berdel and Moryś 2000), and further changes in PV-IN features, including perineuronal nets in the medial prefrontal cortex and BLA, continue through adolescence (Baker et al. 2017). These continuous changes from P10 until adulthood may explain the gradual changes in inhibitory transmission onto BLA PNs across the ages we examined. How other interneuron subtypes, such as somatostatin-and vasoactive intestinal peptide-expressing interneurons, integrate into the BLA circuit, whether their activity changes across development, and how they shape BLA PN connectivity and activity remain largely unknown.

In addition to synaptic changes, several metrics of BLA PN excitability decreased from P11 to P35 and P70. BLA PN cellular development continues over the first weeks of life, with prominent growth of dendritic arbors and soma size (Ryan et al. 2016), consistent with the increase in capacitance we measured. In the following weeks, numerous ionic currents change, including those regulating Ih and the afterhyperpolarization (Ehrlich et al. 2012; Ryan et al. 2016). These changes mirror our observations and likely underlie them.

We also observed cell type-specific changes in intrinsic excitability in PL neurons between P10-11 and P70. As in the BLA, PL PN excitability decreased, whereas excitability of both SST-INs and PV-INs increased across the same period. Parvalbumin is a calcium-binding protein that regulates excitability and plasticity (Orduz et al. 2013) and is strongly developmentally regulated, with low expression in juveniles that increases into adolescence and adulthood (Berchtold and Means 1985; Solbach and Celio 1991; Soriano et al. 1992; Alcántara et al. 1993; del Río et al. 1994; Caballero et al. 2014), when mature spiking properties emerge. In the anterior cingulate cortex, SST-INs show marked developmental changes in intrinsic properties, including maximum firing frequency, with different properties maturing at distinct times (Pan et al. 2016). Prefrontal SST-IN and PV-IN properties mature on different timescales, differing by a couple of days to nearly a week (Yang et al. 2014; Pan et al. 2016). How these trajectories shape the synaptic development of mPFC microcircuits remains incompletely understood. Because different cell classes, such as SST-INs, play distinct roles in fear processing during development than in adulthood, linking the maturation of their activity profiles to fear behavior is a clear next step.

### Implications and next steps

These data position the maturation of prefrontal-amygdala circuitry as a candidate substrate for the emergence of persistent cue-specific fear and define the ages at which it unfolds. The next step is to identify the molecular mechanisms that time mPFC-BLA connectivity and memory persistence, and to ask whether they also underlie the effects of early-life adversity, which accelerates adult-like fear retention in maternally separated rats (Callaghan and Richardson 2012), accelerates the emergence of amygdala-prefrontal connectivity in children (Gee et al. 2013), and alters fear expression and mPFC-BLA connectivity in mice (Manzano Nieves et al. 2020).

## MATERIALS AND METHODS

### Animals

Male and female C57BL/6J mice (Jackson Labs strain # 000664) were used. Behavioral experiments were performed in early juvenile (P14-15), late juvenile (P20-21) and adult (P59-70) mice. For tracing, stereotaxic surgeries were performed at P1, P28 and P80 and tissue was collected at P11, P38 and P90. For optogenetics-assisted electrophysiology, surgeries were performed at P1, P25 and P60 and recordings made at P11, P35 and P70. Other recordings were made at P10-11, P32-36 and P70. SST-IRES-Flp (Jackson Labs strain # 031629) mice were crossed to Ai65F (Jackson Labs strain # 32864) reporter mice to drive specific expression of tdTomato in SST-INs. PV-IRES-Cre (Jackson Labs strain # 17320) mice were crossed to Ai9 (Jackson Labs strain # 007909) reporter mice to drive specific expression of tdTomato in PV-INs. Pups were kept with the dam until weaning at P21, and mice were thereafter group housed (2-6 per cage) on a 12 h light-dark cycle with ad libitum access to food and water. All procedures were approved by the Institutional Animal Care and Use Committee at the Heersink School of Medicine, University of Alabama at Birmingham (APN # 22450 and 22453).

### Cued fear conditioning

Conditioning was performed in sound-attenuating chambers consisting of foam-insulated boxes, house fan, overhead house light, and behavioral arenas. For conditioning, mice were placed in the training context (context A: metal grid floors, square metal walls, and 70% ethanol odor a pan below the floor grid) and allowed a 200 s baseline, followed by six pairings of a tone (conditioned stimulus, CS; 80 dB, 2 kHz, 20 s) with a co-terminating scrambled foot shock (unconditioned stimulus, US; 0.6 mA, 2 s) separated by 80 s intervals. Memory was tested 1, 7 and 14 d after training in the same mice, in a different context (context B: white plastic floor, rounded white plastic walls, and 70% isopropanol odor in a pan below the floor), with a 100 s baseline followed by four 20 s CS presentations without shock separated by 55 s. Freezing, defined as the absence of all movement except respiration for at least 3 s, was scored automatically with VideoFreeze software (MedAssociates) and verified by an experimenter blind to group. Because immobility in young pups may include behaviors other than freezing, we refer to immobility when neither tone nor shock was present.

Cue-evoked freezing was calculated as the mean percent time freezing across the four CS intervals, and baseline freezing as the percent time freezing during the pre-tone period.

### Stereotaxic surgeries

Mice were anesthetized by hypothermia (P1) or isoflurane (P25 and older) and placed in a 3D-printed mold (P1) or a stereotaxic frame. P1 pups were placed on ice until movement stopped (~5 min) and then transferred to a 3D-printed mold, where their bodies and heads were secured with clay. Older mice were placed in a stereotaxic frame once anesthetized. After surgery, mice of all ages were recovered on a heating pad and returned to their home cage with their littermates or cage mates together. Pups were placed in a plastic box containing bedding from their home cage on a pad set to 37-38 °C and were monitored for ~45 min after the last pup in the litter was injected, until their color returned and they moved normally. All pups from a litter were removed from the dam at the same time and returned together. Older mice were similarly monitored on a heating pad until recovery from anesthesia.

AAV1-CaMKIIα-hChR2(H134R)-eYFP (Addgene #26969-AAV1; ≥ 1×10^13^ vg/mL) was bilaterally injected into the mPFC for tracing, or into the PL for electrophysiology. Volumes and coordinates were adjusted at each age to produce comparable spread. For tracing, injections were made at P1 (200 nL per hemisphere at 50 nL/min; 3.15 mm A/P, ±0.2 mm M/L, −0.45 mm D/relative to lambda), P28 (375 nL per hemisphere at 100 nL/min; 1.75 mm A/P, ±0.3 mm M/L, −1.55 mm D/relative to bregma) and P80 (400 nL per hemisphere at 100 nL/min; 1.85 mm A/P, ±0.3 mm M/L, −1.6 mm D/V relative to bregma). For electrophysiology, mice were injected in the PL at P1 (75 nL per hemisphere at 35 nL/min; 3.15 mm A/P, ±0.2 mm M/L, −0.35 mm D/relative to lambda), P25 (200 nL per hemisphere at 100 nL/min; +1.70 mm A/P, ±0.35 mm M/L, −1.4 mm D/V relative to bregma) and P60 (250 nL per hemisphere at 100 nL/min; +1.70 mm A/P, ±0.35 mm M/L, −1.7 mm D/V relative to bregma). Viral expression was allowed to proceed for 10 d before tissue collection or recording (P11, P38 and P90 for tracing; P11, P35 and P70 for electrophysiology).

For tracing, mice were anesthetized and transcardially perfused with ice-cold 1× PBS followed by 4% paraformaldehyde (PFA). Brains were removed, post-fixed overnight in 4% PFA at 4 °C, washed twice in 1× PBS, and stored in 1× PBS at 4 °C. Only mice with expression across most of the mPFC (>70% of mPFC area expressing eYFP) were imaged and quantified. Mice without sufficient mPFC expression were not imaged in the BLA.

### Immunohistochemistry, imaging and quantification

The mPFC and BLA were sectioned at 50 µm at room temperature in PBS on a Leica VT1000S vibratome. Floating sections were processed for immunohistochemistry to amplify eYFP. Sections were blocked in 5% normal donkey serum in PBS with 0.3% Tween 20 for 1 h at room temperature, incubated overnight at 4 °C with chicken anti-GFP (1:500; Abcam #AB16901) in blocking solution, and then incubated for 2 h with FITC-conjugated donkey anti-chicken secondary antibody (1:500; Jackson ImmunoResearch #703-095-155). Sections were stained with DAPI (Sigma-Aldrich #D9542-10MG), mounted on slides (Fisher Scientific #12-550-15), and cover slipped with Fluoro-Gel II mounting medium (Fisher Scientific #50-246-93). Images were acquired on a Keyence BZ-X800 microscope at 10× magnification, and tiles were stitched with the Keyence analysis software. All ages were imaged with identical acquisition settings, and any brightness or contrast adjustments were applied identically across ages. The BLA and mPFC were identified at matched anterior-posterior levels. Infection was quantified as the percentage of mPFC area expressing eYFP. The BLA region of interest was drawn on an overlay created from brain atlas images, and BLA eYFP intensity was measured within it. After background subtraction, the intensity was normalized to the percentage of mPFC area expressing eYFP in the same animal. Only mice with expression across most of the mPFC (>70%; see Stereotaxic surgeries) were imaged in the BLA and quantified. Labeling in the mediodorsal and ventromedial thalamus was assessed qualitatively and not quantified.

### Electrophysiology

Mice were deeply anesthetized with isoflurane before decapitation and brain removal. Acute coronal brain slices (300 µm) containing the BLA or PL were prepared on a Leica VT1200S vibratome in ice-cold, carbogen-bubbled (95% O2, 5% CO2) sucrose-based cutting solution containing (in mM) 210 sucrose, 11 glucose, 26.2 NaHCO3, 2.5 KCl, 1 NaH2PO4, 4 MgCl2, 0.5 CaCl2 and 0.5 Na-ascorbate. Slices recovered for 45 min at 34 °C in carbogen-bubbled artificial cerebrospinal fluid (ACSF) containing (in mM) 119 NaCl, 11 glucose, 26.2 NaHCO3, 2.5 KCl, 1 NaH2PO4, 2 MgCl2 and 2 CaCl2, and were kept at room temperature for the duration of recordings.

Whole-cell recordings used fire-polished borosilicate glass electrodes (3-4 MΩ) pulled on a Narishige PC-100 puller. For voltage-clamp experiments, electrodes contained (in mM) 130 Cs-methanesulfonate, 10 HEPES, 10 Na-phosphocreatine, 1 QX-314, 8 NaCl, 0.5 EGTA, 4 Mg-ATP and 0.4 Na2-GTP (pH 7.25; 298 mOsm). For current-clamp experiments, electrodes contained (in mM) 127.5 K-methanesulfonate, 10 HEPES, 0.6 EGTA, 5 KCl, 2 Mg-ATP, 0.3 Na2-GTP, 5 Na-phosphocreatine and 2 MgCl2 (pH 7.25; 296 mOsm). Cells were visualized with differential interference contrast optics. PL and BLA principal neurons (PNs) were identified by their large pyramidal soma and prominent apical dendrites, and confirmed by their membrane properties and spike waveform. PL PV-INs and SST-INs were identified and patched via expression of tdTomato fluorescence.

Miniature currents were recorded with 1 µM tetrodotoxin in the ACSF. Cells were voltage-clamped at −60 m (mEPSCs) or 0 mV (mIPSCs) and recorded for 3-5 min, with both measured in the same cells. For optogenetically evoked EPSCs, 1 ms, 473 nm light pulses (10-15 mW) were delivered through an LED-coupled microscope objective to stimulate ChR2-expressing axon terminals from the PL, in the presence of 1 µM tetrodotoxin and 100 µM 4-aminopyridine. Paired-pulse recordings used two light pulses at interstimulus intervals of 50 ms, 100 ms, 250 ms, 500 ms and 1 s (~10 sweeps per interval), and paired-pulse ratio was calculated by dividing the amplitude of the second response by the amplitude of the first response. Intrinsic excitability recordings were made in ACSF with 10 µM CNQX, 75 µM AP5 and 100 µM picrotoxin. Cells were held in current clamp at −70 mV via continuous current injection, and 500 ms current steps were applied from-15 pA in 10 pA increments. Passive and active membrane properties were measured. Rheobase is defined as the minimum current that elicited one action potential. Data were acquired at 10 kHz and low-pass filtered at 3 kHz with a Multiclamp 700B amplifier. Recordings were analyzed by experimenters blind to group with Easy Electrophysiology (v2.8.0) or Clampfit 11.3 (Molecular Devices). Recordings were excluded if access resistance changed by more than 20% or if the leak current was unstable or exceeded 200 pA. For BLA optogenetics-assisted recordings, viral targeting in PL was verified by fixing the frontal cortex of recorded mice in 4% PFA overnight, followed by subsequent histological sectioning and fluorescence imaging.

### Statistical analysis

Freezing during training was analyzed within each age group by one-way repeated-measures ANOVA with the Geisser-Greenhouse correction and Dunnett’s multiple comparisons test against baseline. Retrieval at each time points was analyzed by two-way repeated-measures ANOVA (stimulus, matched; age) with Šídák’s multiple comparisons test of baseline vs. CS within each age. CS freezing and CS − baseline differences were compared across ages at each delay by one-way ANOVA with Tukey’s test. For tracing, infection and normalized BLA signal were compared across ages by one-way ANOVA with Tukey’s test. Analyses were performed in GraphPad Prism.

For electrophysiology, normality and homogeneity of variance were tested with Shapiro-Wilk and Levene’s tests, and non-parametric equivalents were used when assumptions were not met. Action-potential input-output curves and paired-pulse ratios were analyzed by two-way repeated-measures ANOVA. Passive membrane properties and miniature current frequency and amplitude were analyzed by one-way ANOVA or the Kruskal-Wallis test, with Tukey’s or Dunn’s post hoc tests, respectively. Two-group comparisons used the Mann-Whitney U-test or an unpaired t-test. Cells were treated as the unit of analysis. Analyses were performed in OriginPro 2025b SR1 10.2.5.234.

Experimenters were blind to group wherever possible, including for electrophysiology analysis, behavioral scoring, and image quantification; blinding to age was not possible during behavioral testing because pups of different ages differ in appearance.

## AUTHOR CONTRIBUTIONS

**Verdion Martina** Conceptualization, Data curation, Formal analysis, Investigation, Methodology, Project administration, Validation, Visualization, Writing - original draft, Writing - review & editing.

**Zephyr R. Desa** Data curation, Formal analysis, Funding acquisition, Investigation, Methodology, Validation, Visualization, Writing - original draft, Writing - review & editing.

**Rodrigo Campos-Cardoso** Data curation, Formal analysis, Investigation, Methodology, and Validation.

**Terry Lewis** Investigation, Methodology, Project administration, Supervision

**HaoSheng Sun** Conceptualization, Data curation, Formal analysis, Funding acquisition, Project administration, Supervision, Validation, Visualization, Writing - original draft, Writing - review & editing. **Kirstie A. Cummings** Conceptualization, Data curation, Formal analysis, Funding acquisition, Project administration, Supervision, Validation, Visualization, Writing - original draft, Writing - review & editing.

